# Unraveling the Role of Azobenzene-Based Photoswitchable Lipids in Controlling Innate Immune Response

**DOI:** 10.1101/2024.07.28.605494

**Authors:** Anastasiia Delova, Aline Makhloutah, Yann Bernhard, Andreea Pasc, Antonio Monari

## Abstract

The control of the activation of the innate immune response, notably by using suitable visible light sources, may lead the way to the development of phototimmunotherapeutic strategies. In this contribution we analyze the effects of the E/Z interconversion on a phosphatidyl serine lipid containing an isomerizable azobenzene moiety, which has recently been shown to regulate the activation of natural killer cells and the production of cytokines [*J. Am. Chem. Soc.* 2022, 144, 3863−3874]. In particular we analyze the differential interactions which the innate immune system TIM-3 sensors. We show, resorting to long-range molecular dynamic simulations including enhanced sampling, that the Z isomer lead to a slight decrease of the binding free energy coupled with a less pronounced rigidification of the protein compared to the E isomer and the native lipid, justifying its less pronounced activation of the immune response.

TOC GRAPHIC

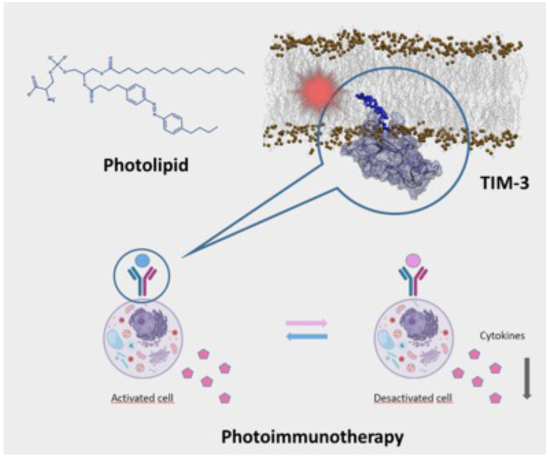

## INTRODUCTION

The innate immune serves as the front line of the immune response against the attacks of exogenous pathogens or endogenous agents affecting the viability of cells and organisms.^1,2^ Therefore, maintaining its proper regulation is crucial, while its aberrant behavior is usually connected to a number of seriously debilitating pathologies, including as a most paradigmatic examples autoimmune disease.^3^ Usually the activation of the innate response triggers apoptotic signaling, which leads to the cell death thus halting the propagation of the infections.^4^ Furthermore, the stimulation of interferon and cytokines production correlates with a proinflammatory response which induces an unfavorable environment for the pathogens.^5,6^ Different innate immune system pathways exist, which may also respond to different threats, such as the OAS/RNAse L pathways,^7–9^ which is tailored to sensing the presence of exogenous RNA, or the cGAS-STING^10,11^ which is instead most sensitive to the presence of exogenous DNA. Recently, the role of the T cell transmembrane, Immunoglobulin, and Mucin (TIM) proteins has been particularly recognized for its diverse effects in regulating the innate immune response. TIM proteins exist in different isoforms^12–14^ with TIM-1, TIM-2, TIM-3, and TIM-4 being the most widely studied. They are large transmembrane proteins (Figure 1) comprising different domain, including a cytosolic region, a transmembrane segment, a linker mucin domain involving glycosylation sites,^13^ and finally the N-terminal immunoglobulin–like (IgV) domain. In the case of TIM-1, TIM-3, and TIM-4 the IgV domain senses the external lipid leaflet of other cellular membranes (Figure 1B) and interacts specifically with lipids whose polar heads are comprised of phosphatidyl-serine (PDS) moieties.^12,15–19^ This specificity is at the base of its action since it characterizes them as efficient sensors of apoptotic cells.^20–28^ Indeed, during apoptosis PDS-containing lipids migrate from the inner to the outer leaflet of the cellular membrane.^22,27^

**Figure 1.**
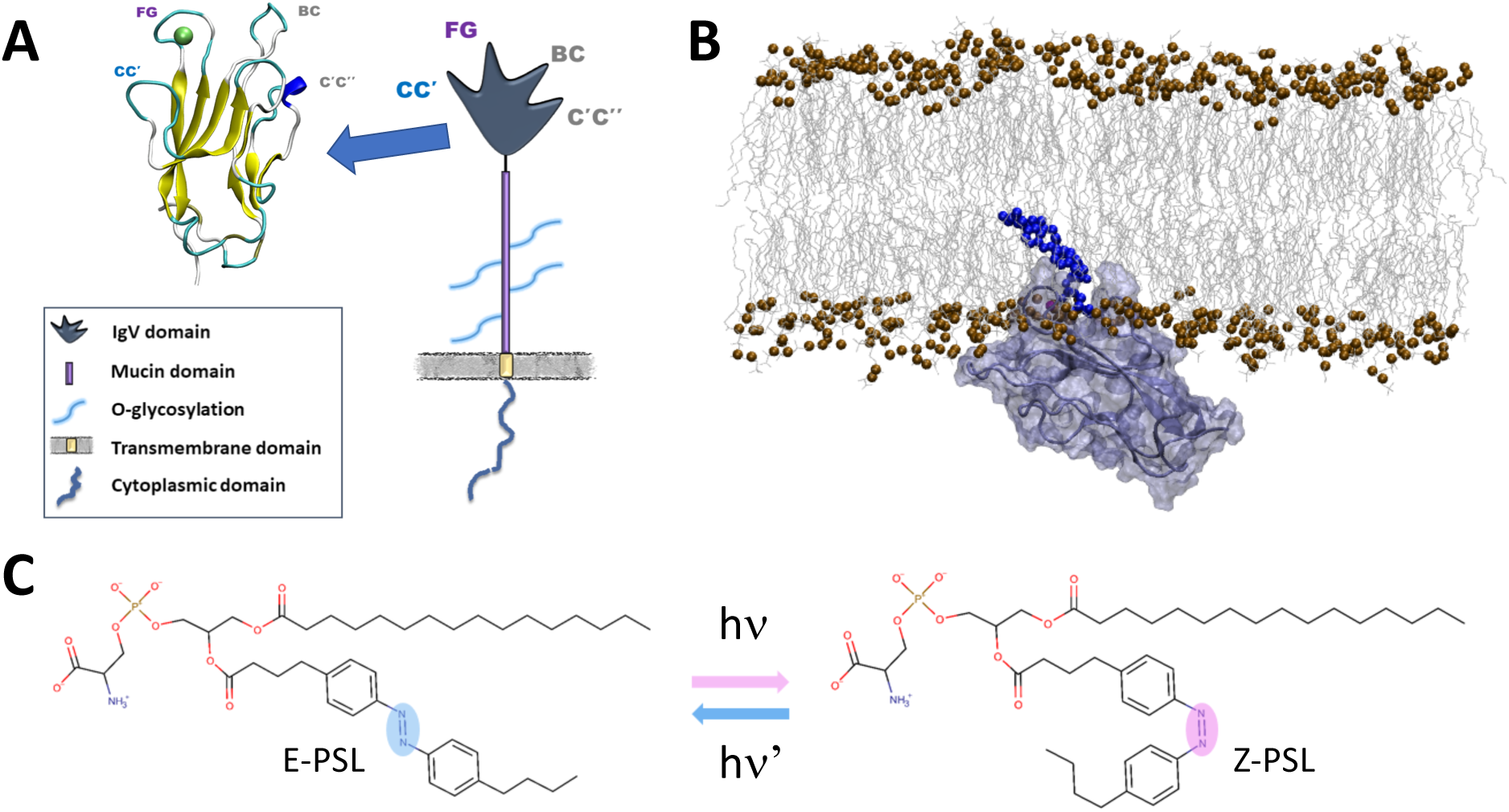
Schematic structure of TIM-3 highlighting the IvG domain at atomic recognition (A). Snapshot extracted from the MD simulation and showing the interaction of the IvG domain with a lipid membrane and its binding to a PDS-containing lipid (B). Representation of the E and Z isomers od the photoactivable lipids used in this study (C).

In contrast with the other members of the family, TIM-2 is not interacting with PDS and should, thus, be related to a different mechanism of actions, which remains elusive to date.^29–31^ Yet the role of TIM is more complex and is not restricted to the simple recognition of apoptotic cells. Indeed, a complex balance between the activation of anti-inflammatory signals, which helps in maintaining immune response tolerance, and the production of cytokine has been observed, pointing to a multifaced and highly dynamic role.^32^

TIM-3, which is preferentially expressed in T-helper 1 and T-cells 1,^33,34^ but also found in dendritic cells,^35^ is particularly standing out among the different members of the family. TIM-3 also represents a marker of the maturation of natural killer (NK) cells,^6,29^ while its activation correlates with cytokine production^36,37^ and is related to autoimmune diseases.^38^ More recently, the role of TIM-3 in regulating the tumor microenvironment has been particularly pointed out.^39–42^ Indeed, TIM-3 is also related to the exhaustion of T-cells in tumors or to the functionality loss of lymphocytes in some tumor microenvironments. Indeed, the overexpression of TIM-3 may be considered as a cancer biomarker, correlating with a bad prognosis,^43–45^ while the blockade of TIM-3 activity has shown beneficial effects, even for the treatment of tumors in advanced stages.^46–48^ Structurally the IvG domain of TIM-3 is composed of a rigid core, rich in β barrel motifs, flanked by four flexible loops. Notably the pocket between the FG and CC’ loops constitute the PDS binding site (Figure 1A).^49,50^ The interaction with the phosphatidyl serine is mediated by a Ca^2+^ ion and by persistent hydrogen bonds with different protein residues, notably D102, Q43, and S42.^49^ Recently, by using enhanced sampling and extended Molecular Dynamics (MD) simulations the thermodynamic parameters of the interaction between the IvG domain and PDS containing lipids have been determined.^16^ Notably, we have shown that the inclusion of the lipid in the binding pocket is barrierless and favored by a free energy driving force of about 20 kcal/mol.^16^ While the polar head is held tight between the FG and CC’ loops the lipid tails remains globally parallel to the membrane axis and stretch away from the IvG surface.

It has been shown that PDS-containing photoswitchable lipids (PSL) including an azobenzene moiety in one of the lipid tails (Figure 1C) may act as regulator of the activity of TIM-3.^51^ Indeed, under irradiation of suitable UV/Vis light azobenzene experiences a fast E/Z photoisomerization^52,53^ leading to the conversion of the E-PSL to the Z-PSL isomer. While the former is comparable to the native PDS lipids concerning the induction of TIM-3 response, the latter is reducing the activation of the protein.^51^ These results pave the way to the exploitation of molecular photoswitches to enforce the light-controlled activation of the immune system response. Due, to the involvement of TIM-3 in cancer development and in shaping the tumor microenvironment, this strategy may be highly suitable for photoimmunotherapy purposes. However, the molecular bases leading to the differential activity of the two isomers, and hence the different activation of the protein, remains up to date elusive precluding the design of improved photoimmunotherapy agents.

In this contribution, by using MD simulations and enhanced sampling protocols, we explore the interaction of the two PSL isomers with the TIM-3 IvG domain, in presence of a realistic lipid bilayer to mimic physiologically relevant conditions. In particular we show that the Z isomers presents a slightly less favorable binding free energy with TIM-3 than the E-isomer, while it also induces a less pronounced rigidification of the protein scaffold, globally justifying the reduced activation of the immune response.

## COMPUTATIONAL METHODOLOGY

The initial structures of the PSL in E- and Z-conformations have been drawn using the Marvin Sketch chemical drawing software.^54^ The structures of both isomers have been optimized at the B3LYP/6-31G(d) density functional theory, considering an implicit water solvent modeled with the PCM approach using the Gaussian09 software^55^. On top of the optimized geometry we have calculated the restrained electrostatic potential (RESP) charges using the HF/6-31G* level of theory. Then the antechamber utilities of AmberTools22^56^ package were used to generate Generalized Amber Force Field (GAFF) parameters for both conformers. To precisely model the azobenzene unit, and especially the torsion along the isomerizable double bond, we have replaced the original amber parameters with those described by S. Osella et al.^57^

The CHARMM-GUI^58^ web-interface was used to build an initial model of a lipid membrane bilayer having a composition characteristic of normal eukaryotic cells. A mix of lipids, including POPC, POPE, POPS, and cholesterol, was used with respective molar ratios of 30%, 25%, 15%, and 30%. This resulted in a total of 200 lipids in each leaflet coherently with the strategy used in our previous work^59^. A 30 Å water buffer containing a physiological concentration of 0.15 M of K^+^ and Cl^-^ ions was also included and added to the system. The IgV domain of the mice TIM-3 protein (PDB: 3KAA)^60^ has been chosen as the starting structure. The protein was placed in close proximity to the lipid’s polar heads within the bulk water to allow the interactions with the lipids. Lipids have been represented by Amber Lipid14 force field^61^, while water was described using TIP3P^62,63^ and the protein with the Amber ff14SB^64^ protein force field.

To place one PSL lipid in the IvG binding site we used a strategy similar to the one of our previous work:^59^ namely, we considered the IvG system interacting with a PDS lipid which has been obtained from a 1 μs MD simulation. Since the polar head (phosphatidyl-serine) of the native PDS lipid is identical to the one of PSL, we aligned the latter, either in E-or Z-conformation, with the native PDS lipid using the VMD program to obtain starting conformations for E-or Z-PSL.

For both initial systems we performed all-atom classical MD simulations using NAMD^65,66^ software, followed by analysis and visualization with the VMD program^67^ Hydrogen mass repartition^68^ (HMR) was employed allowing the use of a 3.0 fs time-step in the propagation of the simulation via the numerical resolution of Newton equations of motion, also using Rattle and Shake algorithms.^69^ Note that despite longer equilibration the use of 4 fs time-step, which should be compatible with HMR resulted in numerical instabilities and was, thus, discarded. For all the systems the minimization, thermalization and equilibration steps were performed, by gradually removing positional constraints on non-water heavy atoms over a 12 ns period. Throughout equilibration and production, a constant temperature of 300 K and a pressure of 1 atm have been maintained, leading to a isothermal and isobaric (NPT) ensemble in which the lipid membrane remains in its liquid phase. The Langevin thermostat^70^ and barostat^71^ were used to enforce temperature and pressure conservation, respectively. The production phase for both systems spanned a total simulation time of 1 μs. To analyze the stability of the E/Z-PSL-protein complex the root mean square deviation (RMSD)^72^ and root mean square fluctuation (RMSF)^73^ were calculated.

Umbrella Sampling (US)^74^ was used to obtain a potential of mean force (PMF) describing the interaction between the PSL’s polar head and the binding pocket comprised between the FG and the CC’ loop of the IgV domain. As a collective variable for US we choose the distance from the center of mass of the bound PSL’s polar head to the center of mass of the binding pocket, which includes Ca^2+^ and the four principal interacting residues: S42, Q43, Q95, and D102.A pictorial representation of the collective variable is also provided in SI. The collective variable value was increased from 3.0 to 28.0 Å with consecutive windows separated by a step of of 0.5 Å. Each US window has been propagated for 180 ns for the E-PSL and 240 ns for the Z-isomer to ensure a good equilibration with a force of 22.5 kcal/(mol.Å).

The final merged PMF profile was generated by integrating the results from the different windows using the Weighted Histogram Analysis Method (WHAM)^75^. The overlap of the distribution of the collective variables among the different windows can be appreciated in SI, as well as the analysis of the convergence of the PMF. Linear Interaction Energy (LIE)^76^ was used to analyze the contribution of PSL to the binding with the TIM-3 protein. LIE calculations were performed for both the E- and Z-PSL isomers, considering the full structure, which includes the hydrophobic chains. The calculations accounted for the electrostatic (EELEC) and van der Waals (EVDW) interaction energies. In addition to US, Molecular Mechanics Poisson-Boltzmann Surface Area (MMPBSA)^77^ approach has been used to estimate the binding free energy contributions of the individual residues involved in the interaction between the PSL and TIM-3. Per-residue decomposition analysis was performed to determine the energetic contributions of specific residues located within the FG CC’ loop (residues 42-44, 95-104), as well as Ca^2+^. MMPBSA calculations were performed on top of the equilibrated MD simulations using the Amber interface.

## RESULTS AND DISCUSSION

MD trajectories exceeding the μs time-scale have been propagated for PSL interacting with IvG. Both the E- and Z-PSL isomers have shown the formation of a persistent bound-states with the IvG domain, consistently remaining inside the binding pocket of the protein (Figure 2). The hydrophobic tails of the E isomer behave similarly to those of native lipids^16^ and extend towards the hydrophobic core of the membrane (Figure 2A). Conversely, Z-PSL adopts a more globular arrangement, which develops more pronounced interactions between the lipid tails and the protein loops, which may also lead to partial steric clashes (Figure 2B).

**Figure 2.**
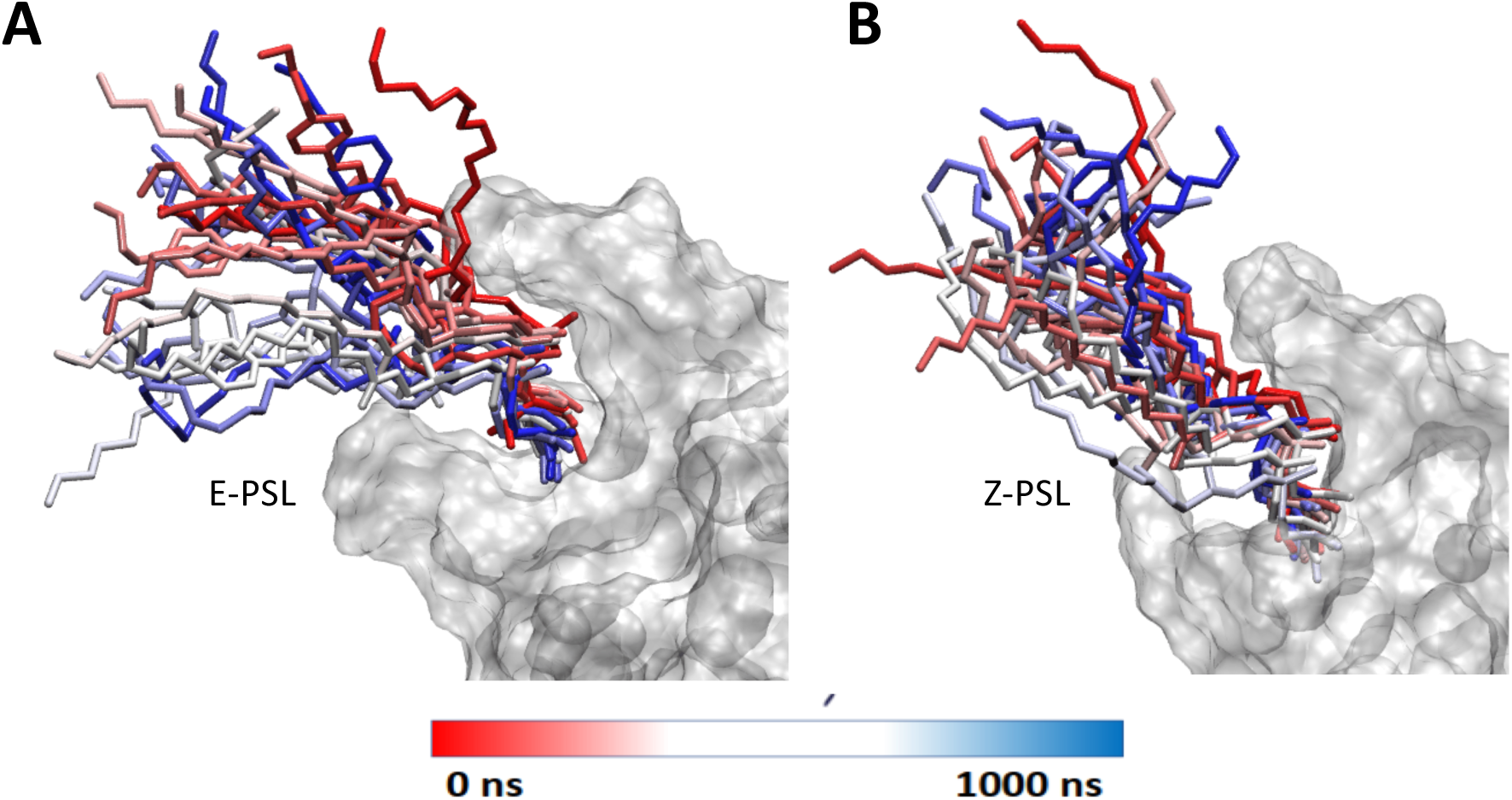
Snapshots showing the time evolution of the positioning of the E- and Z-PSL (panel A and B, respectively) inside the binding pocket of the IvG domain. The protein is represented as a transparent surface, while the PSL is in licorice.

To better quantify the persistence of the interactions locking PSL into the protein binding pocket we have analyzed the evolution of the main interactions between the protein residues and the lipid polar head as shown in Figure 3 and Table 1. These results are also compared to the experimental structure obtained for an isolated PDS polar head interacting with the IvG domain.^18^

**Figure 3.**
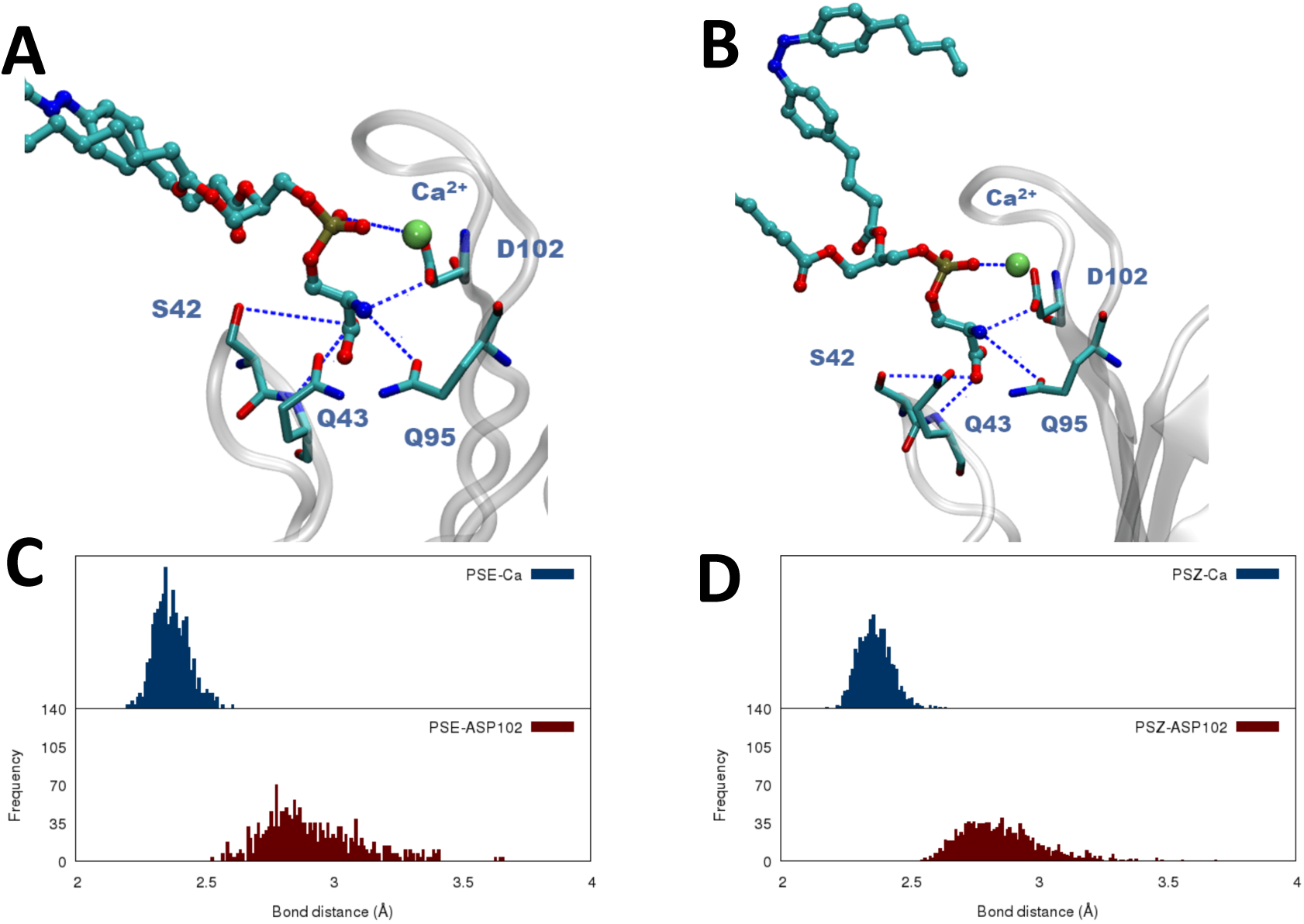
Representative snapshots showing the disposition of the E-PSL (A) and Z-PSL (B) in the IvG binding pocket. Distribution of the main distances along the MD simulation for the E-(C) and Z isomer (D), respectively. This behavior, as well as the persistence of the interaction in the IvG binding pocket can also be appreciated in Figure 3, in which we also point out the strong interaction with Ca^2+^ which is characterized by a very peaked distribution with a maximum at short distances. Interestingly, once again these results are coherent with those obtained for the native lipid and reported in a previous contribution.^16^

**Table 1.** Average values and standard deviations of the distances between the PDS unit of E- and Z-PSL and the main IvG residues in the binding pocket. The values for a reference crystal structure of an isolated PDS polar head interacting with IvG are also given as a reference.^18^ The individual atoms used to calculate the distances are represented in Figure 3.

|  | Native PDS<br>(Crystal) | E-PSL | Z-PSL |
| --- | --- | --- | --- |
| Ca<sup>2+</sup> | 2.938 | 2.38 ± 0.06 | 2.37 ± 0.06 |
| Q95 | 3.420 | 3.07 ± 0.42 | 3.62 ± 0.94 |
| D102 | 2.666 | 2.92 ± 0.19 | 2.86 ± 0.16 |
| Q43 | 3.084 | 4.63 ± 0.99 | 4.53 ± 1.09 |
| S42 | 2.750 | 4.94 ± 1.20 | 5.13 ± 1.22 |

From the analysis of the average distances reported in Table 1 no significant differences in the interaction patterns between the two isomers and the native crystal structure may be evidenced, excluding the partial exception of Q95 which shows a slightly larger average distance, as well as a more pronounced dispersion, in the case of the Z isomer. Interestingly, this residue is located in the FG loops, which is the one showing more consistent interactions with the bent lipid tails in the Z isomer. Coherently with our previous results, relative to a native PDS-containing lipid, we may also see that the interaction with S42 is consistently loosened compared to the crystal structure. This effect, which we have interpreted as due to the presence of the lipid tails missing in the reference crystal structure,^18^ is also slightly more pronounced for the Z-isomer in particular considering the dispersion of the distances.

We performed LIE analysis to further quantify the individual energetic contributions stabilizing the E- and Z-isomers in the protein binding pocket. The electrostatic (EELEC) and van der Waals (EVDW) interaction energies between PSL and the protein, including Ca²⁺, reveal that both isomers exhibit favorable binding, albeit with slight differences. The E-isomer displays a mean EELEC of −178 ± 17 kcal/mol and a mean EVDW of −13.96 ± 5 kcal/mol. In contrast, the Z isomer has a slightly weaker electrostatic interaction energy of −172.42 ± 12 kcal/mol and an almost unchanged van der Waals contribution of −13.24 ± 6 kcal/mol (Table 2). These results confirm that electrostatic interactions drive the overall binding of the PSL especially for the E isomer.

**Table 2.** MMPBSA per-residue enthalpy decomposition.

| Residue | Average of total energy decomposition (kcal/mol) |  |
| --- | --- | --- |
|  | E- conformation: | Z-conformation: |
| S42 | -3,07 | -2,15 |
| Q43 | -3,78 | -4,48 |
| C44 | 0,15 | -0,08 |
| Q95 | -2,00 | -0,66 |
| F96 | -0,80 | -0,92 |
| P97 | -0,09 | -0,15 |
| G98 | -0,50 | -0,55 |
| L99 | -4,28 | -3,91 |
| M100 | -2,56 | -3,99 |
| N101 | 0,24 | 0,05 |
| D102 | -5,85 | -3,40 |
| K103 | 0,12 | 0,12 |
| K104 | 0,38 | 0,39 |

Therefore, from these results one would conclude on a rather minor effect of the E/Z isomerization of the PSL in modulating the interaction pathways involving the IvG domain and thus the activation of TIM-3.

However, by analyzing the RMSF plots for the IvG domain interacting either with the Z-or the E-PSL isomer (Figure 4) significant, albeit moderate, differences can be underlined. Indeed, in addition to the N-terminal domain, a noticeable increase of flexibility can be observed for the aminoacids belonging to the FG loop. This concerns in particular the sequence going from residue Q95 to D102, i.e. aminoacids, which are directly located in the binding pocket and participate to the stabilization of the phosphatidyl serine polar head. This change can once again be ascribed to the effect of the lipid tail, which in the Z isomer are susceptible to interact stronger with the FG loop causing partial steric clashes. In turn, this perturbation may be translated into a slightly weakening of the FG/PDS interactions which could ultimately result into an increased flexibility. Note that additional peaks corresponding to either flexibility increase or decrease may be evidenced in Figure 4, however as shown in ESI, they involve residues located either in the N-terminal domain or far apart from the binding site and the lipids. The loss of rigidity of the FG loop, in addition to the weakening of the binding, may also have a secondary effect by partially perturbing the regulation of the signal transduction. This hypothesis is also strengthened by the fact that a secondary region showing a consistent increase of flexibility when interacting with the Z-PSL emerges around residues 80, i.e. in the disordered loop which is not interacting with the membrane and is, in the full TIM-3, interacting with the mucin domain. Conversely, the CC’ (residues 42-45) and the C’C’’ (residues 65-70) loops are slightly less flexible when the IvG domain is interacting with the Z-PSL.

**Figure 4.**
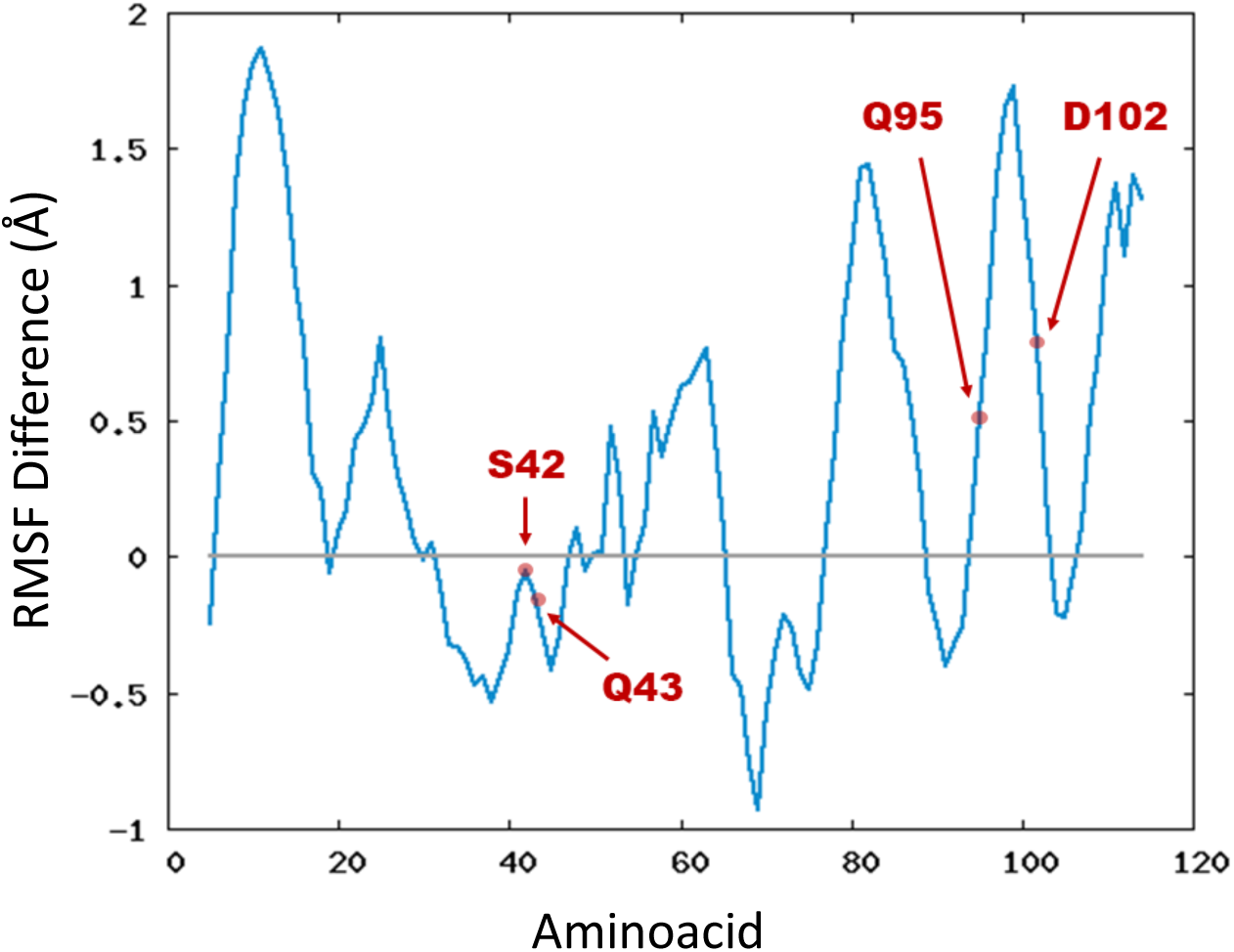
Difference of RMSF of the IvG domain calculated as the RMSF for the protein interacting with the Z-PSL minus the one interacting with the E-PSL. Thus, positive values indicate a greater flexibility for the Z-PSL bound system, while negative ones fro the E-PSL. Individual RMSF plot are provided in ESI. Individual RMSF of the IgV domain in presence of E- and Z-PSL are provided in ESI, along with snapshots showing all the residues presenting significant changes in their RMSF values.

The enthalpic contribution to the binding free energy was also estimated using MMPBSA including per-residue decomposition analysis identifying the role of key protein residues and Ca²⁺. The total binding enthalpy for the E isomer amounts at −128 ± 2 kcal/mol, whereas the Z isomer is slightly less stabilized having a binding enthalpy of −124 ± 1 kcal/mol. Interestingly, the observed small difference in the total binding enthalpy is also consistent with the LIE analysis, strengthening the fact that while both isomers form stable complexes within the IgV domain, the Z isomer’s binding mode is slightly more dynamic due to the increased steric interactions with the FG CC’ loop. The per-residue decomposition of the binding enthalpy for both isomers (Table 2 and ESI) also highlights subtle but significant changes in the interaction pathways, contributing to the observed decrease of the binding affinity for the Z-isomer.

In particular we may note that Q95 contributes more favorably to the binding of the E-isomer (−2.00 kcal/mol) than the Z-isomer (−0.97 kcal/mol), as well as D102 (−5.85 kcal/mol vs. −5.09 kcal/mol for E- and Z-, respectively). Conversely, Q43 exhibits a noticeably stronger interaction with the Z-(−5.52 kcal/mol) compared to the E-isomer (−3.78 kcal/mol), which may reflect a partial reorganization of the hydrogen bond network. In addition, hydrophobic interactions are also slightly altered between the two isomers. Notably, L99, a key residue within the FG loop, contributes more to the binding of the Z-isomer than the E −5.70 kcal/mol vs. −4.28 kcal/mol. Interestingly, Ca²⁺, which plays a critical role in the ligand stabilization, shows a slightly greater interaction energy in the Z-isomer (16.02 kcal/mol) compared to the E-isomer (15.69 kcal/mol). This suggests that minor structural adjustments in the Z-isomer’s binding pose might slightly alter the coordination environment of Ca²⁺, though not to a degree that significantly impacts overall binding affinity. So we show that the E-isomer benefits from a stronger electrostatic stabilization network, particularly through ASP 98 and GLN 91, whereas the Z-isomer gains increased hydrophobic contributions from residues in the FG loop, such as LEU 95.

To comprehensively characterize the thermodynamics of the binding of the two isomers to the IgV domain, we computed, using US, the free energy profile along the distance between the center of mass of PDS and the key residues defining the binding pocket (see ESI for the precise definition of the collective variable). The resulting PMF (Figure 5) presents similar characteristics, indicating that the binding of the lipid occurs without significant energy barriers. Furthermore, both isomers present a minimum at around 5 Å corresponding to the stable bound state. However, the free energy profile of the E isomer is steeper, suggesting that once bound, E-PSL is more rigidly confined within the binding pocket, exploring a more restricted conformational space. Furthermore, the binding free energy of the E-isomer (21.1 kcal/mol) is slightly greater than that of Z-isomer (18.3 kcal/mol). The broader PMF profile obtained for the Z isomer suggests that, likely due to steric interactions involving the lipid tail and azobenzene units, this isomer explores a more extended conformational space and can reach greater distances from the binding site. These findings are consistent with the differences observed in the RMSF profile, which reveals increased flexibility of the IgV domain when bound to the Z-PSL isomer. Although the overall effects of photoisomerization on the interaction patterns between PSL lipids and the IgV domain of TIM-3 are relatively modest, they remain significant, and may correlate with the loss of activity observed upon photoisomerization.

**Figure 5.**
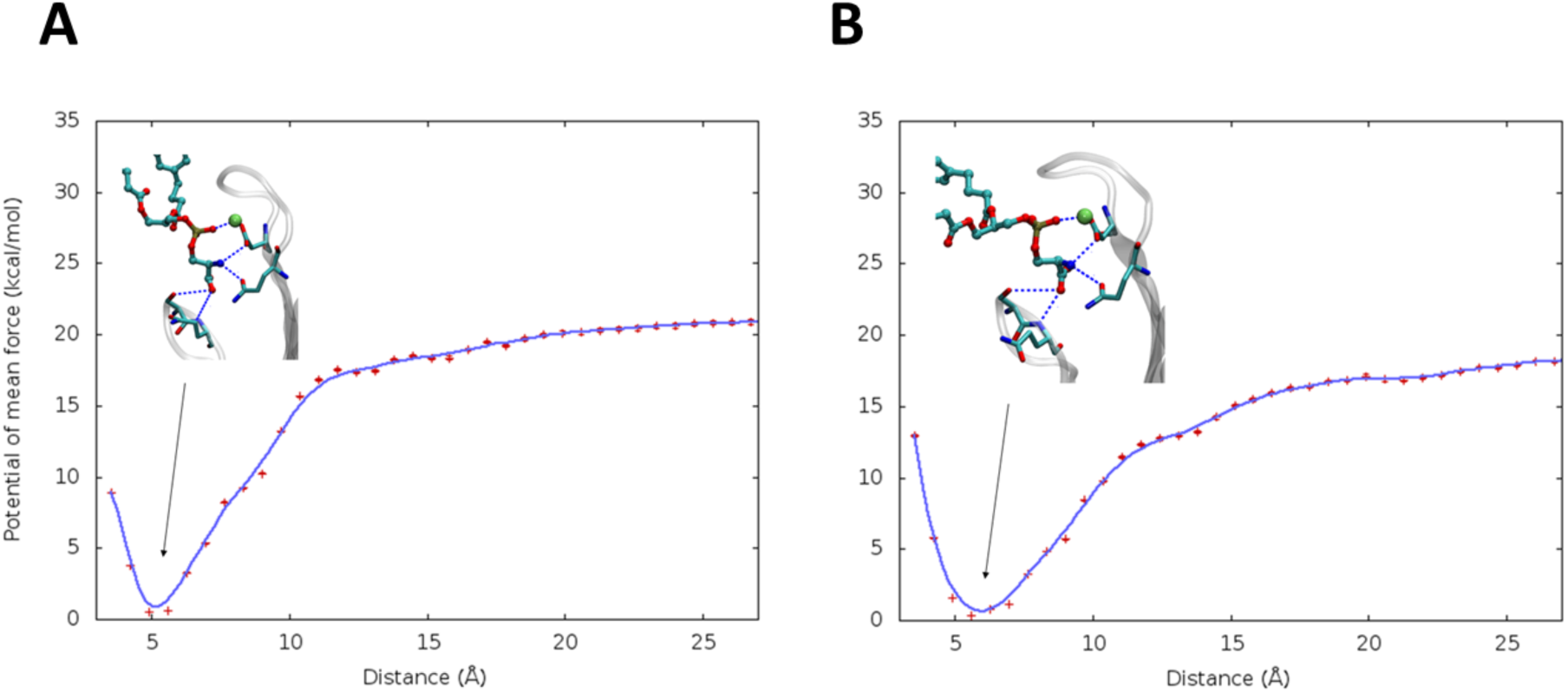
Potential of Mean Force along the collective variable defined as the distance from the binding pocket for the E-(A) and Z-PDS (B) isomers. The respective structures at the equilibrium distance are also given as inlays.

## CONCLUSIONS

We have performed long-range MD simulations of the IvG domain of TIM-3 in presence of a lipid bilayer exceeding the μs range and complemented with enhanced sampling. We have analyzed, at a molecular level, the effect of the photoisomerization of modified lipids including an azobenzene moiety in shaping the activation of the innate immune system TIM-3 agent. In particular we have shown that both isomers interact persistently in the protein binding pocket. Importantly, the interactions patterns are also similar to the ones developed by the native lipid. However, the Z-PSL conformation may induce some partial steric clashes between the lipid tails and the IvG domain of TIM-3. This is in turn translates in a more pronounced flexibility of the loops of the IvG domain, an effect which may also perturb the activation of the protein upon the sensing of phosphatidyl lipids. These global observations are also confirmed by the PMF obtained at the US level. Indeed, the Z-PSL lipid presents a slightly lower binding free energy and a a swallower free energy profile compared to the E-PSL lipid. This observation may suggest that the lower binding free energy, as well as the larger conformational space visited by the Z-PSL lipid and the more important flexibility of the protein correlate with its loss of activity.

Our results establish the molecular bases underlying the regulation of TIM-3 activity by recently proposed photoswitchable lipids which can find applications in the context of photoimmunotherapy. Yet, we also evidence how the perturbation brought by the photoisomerization of the originally proposed azobenzene-based lipids are globally modest, and hence can be non-ideal to provide suitable therapeutic effects. Yet, our work settles the basis for the rational design of improved agents for photoimmunotherapy targeting TIM-3 and its IvG domain, increasing its activity and selectivity. Notably, biomimetic modified lipids showing an improved regulation of TIM-3 will be the subject of a forthcoming contribution.

## ASSOCIATED CONTENT

Time series of the RMSD for the lipid bilayer and the IgV domain, individual RMSF for the IgV domain interacting with the E- and Z-PSL and identification of the most involved residues, interactions between the Ca^2+^ ion and the crucial protein residues, per-residue contribution to the MMPBSA energy, definition of the collective variable, distribution of the collective variable over the different windows, evaluation of the convergence of the PMF calculations for E-PSL and Z-PSL, force field parameters for the isomerizable azobenzene unit. (file type, PDF).

## AUTHOR INFORMATION

### Author Contributions

The manuscript was written through contributions of all authors. All authors have given approval to the final version of the manuscript.

## ACKNOWLEDGMENT

The authors thank GENCI and Explor computing centers and the Platform P3MB for computational resources. A. M. thanks ANR and CGI for their financial support of this work through Labex SEAM ANR 11 LABX 086, ANR 11 IDEX 05 02. The support of the IdEx “Université Paris 2019” ANR-18-IDEX-0001. Support from the Regional Development Fund of the European Union (Programme opérationnel FEDER-FSE Lorraine et Massif des Vosges 2014– 2020/ “Fire Light” project: Photo-bio-active molecules and nanoparticles”) for financial support.

