## Supplementary Information for "Unraveling the Role of Azobenzene-Based Photoswitchable Lipids in Controlling Innate Immune Response"

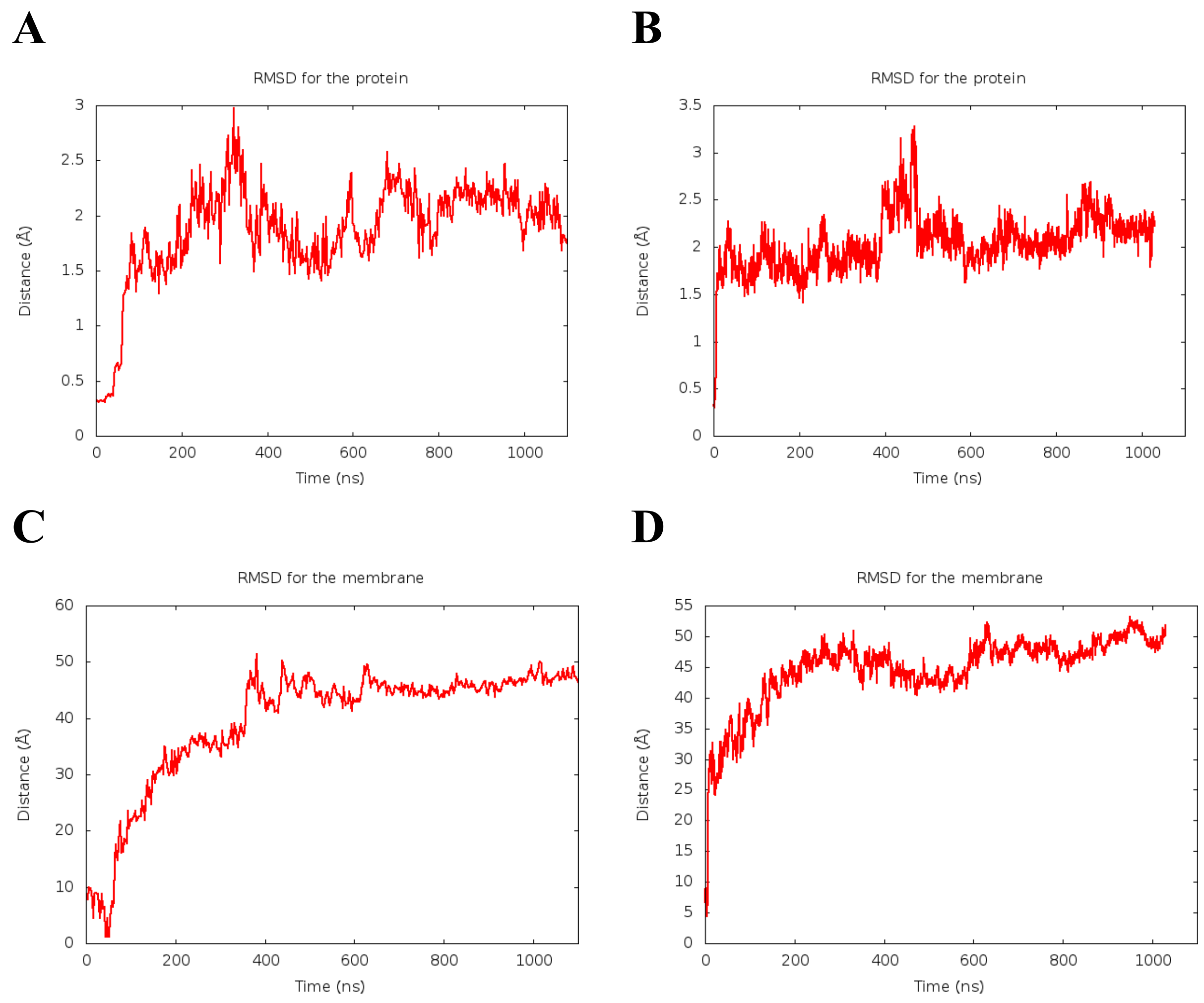


Figure S 1. Time evolution of the RMSD for IvG domain interacting with E-PSL (A) and Z-PSL (B). The RMSD for the lipid bilayer when IvG is interacting with E-PSL (C) and Z-PSL (D) are also provided.

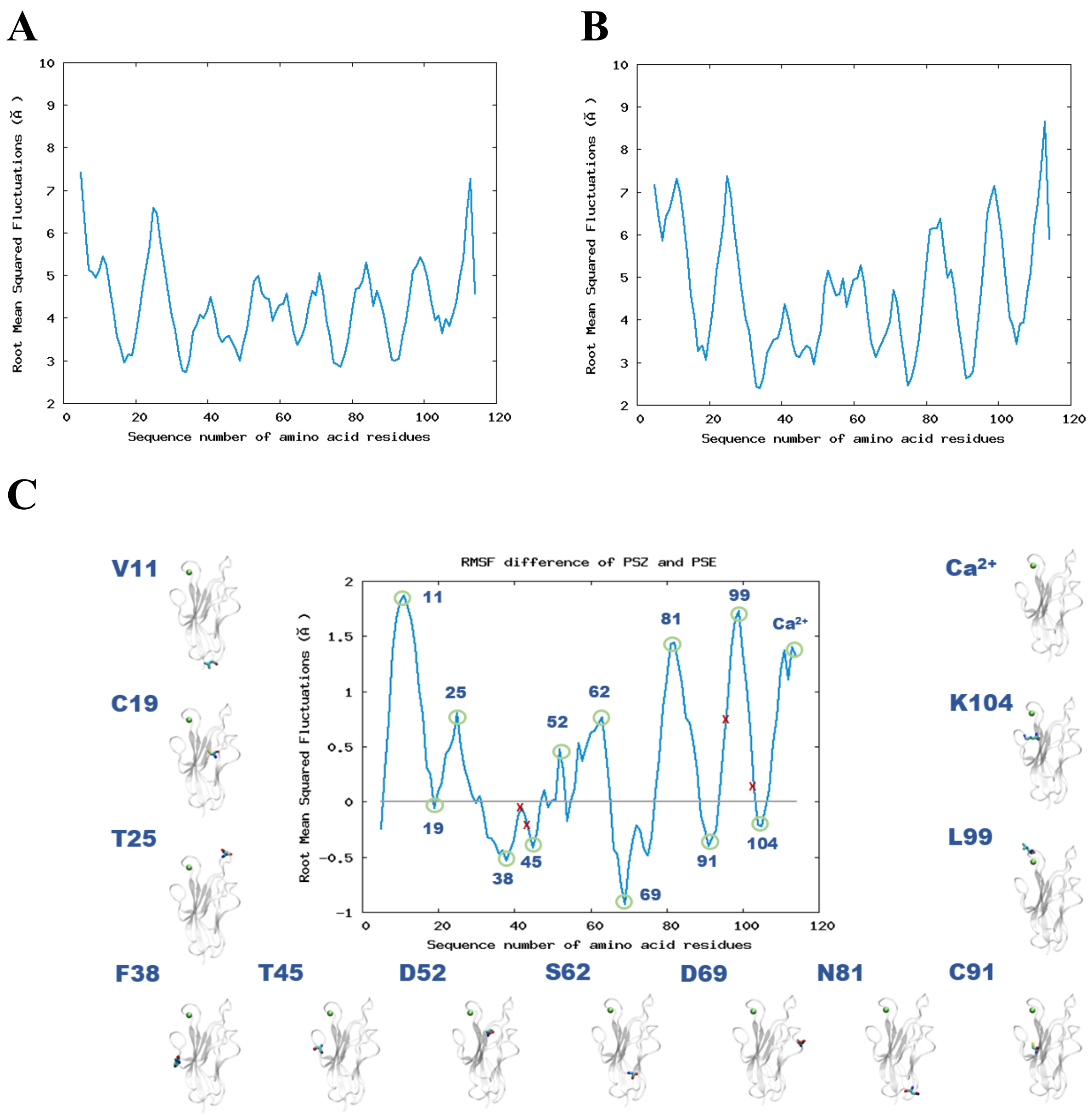


Figure S2. Per-residue RMSF for the IgV domain interacting with E-PSL (A) and Z-PSL (B), as well as the difference in RMSF, calculated as the RMSF of the protein interacting with Z-PSL minus that of the protein interacting with E-PSL (C), are presented. Positive values indicate greater flexibility in the Z-PSL-bound system, while negative values indicate increased flexibility in the E-PSL-bound system. Red crosses mark key protein residues in the binding pockets (S42, Q43, Q95, D102). Snapshots showing the position of residues experiencing the largest changes in RMSF are also provided.

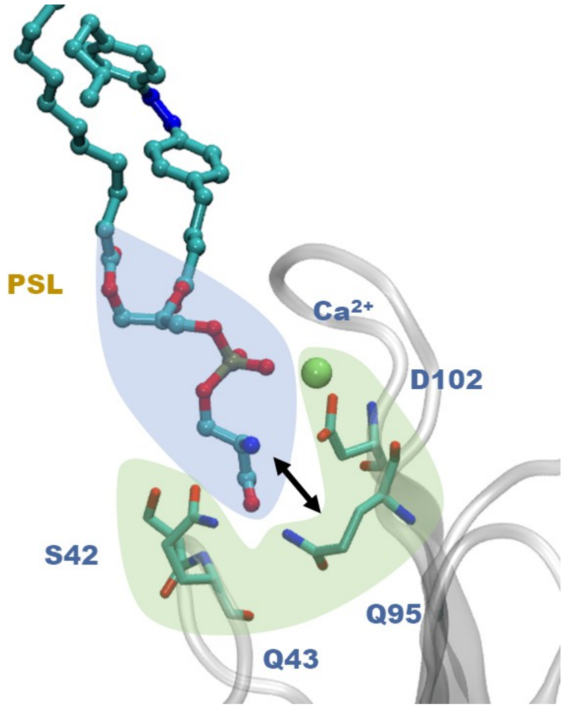


Figure S3. Representation of the collective variable used for the US calculation. The coordinate comprise he distance between the lipid polar heads (the involved atoms are highlighted in purple) and the interaction pocket of the IvG domain (highlighted in green) encompassing the $Ca^{2+}$ ion and the residues S42, Q43, Q95, and D102.

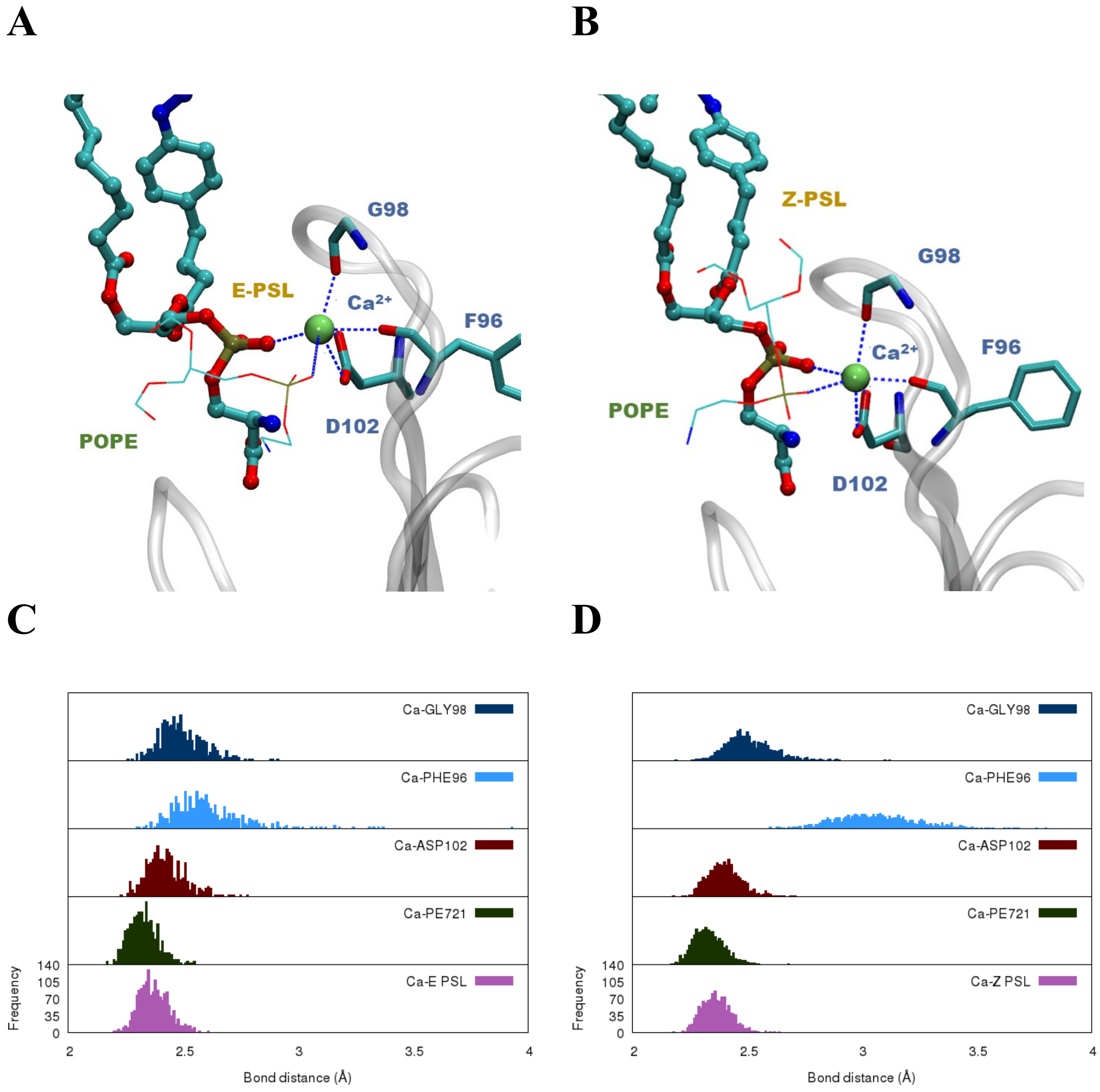


Figure S4. Representative snapshot showing the interactions developed by $Ca^{2+}$ with the IvG residues (F96, G98, D102), PSL molecule (colored in licorice) and POPE lipid (colored in lines mode) for the systems containing E-PSL (A) and Z-PSL (B), respectively. The distribution of the crucial distances between $Ca^{2+}$ and the protein residues are given for E-PSL (C) and Z-PSL (D).

| $Ca^{2+}$ | Native PDS (Crystal) | E-PSL | Z-PSL |
|---|---|---|---|
| PSL | 2.938 | 2.38 ± 0.06 | 2.37 ± 0.06 |
| G98 | 2.711 | 2.50 ± 0.09 | 2.52 ± 0.13 |
| F96 | 2.578 | 2.61 ± 0.17 | 3.06 ± 0.18 |
| D102 | 3.015 | 2.46 ± 0.32 | 2.46 ± 0.11 |
| PE721 | - | 2.33 ± 0.65 | 2.34 ± 0.67 |

Table S1. Average values and standard deviations of the distances between $Ca^{2+}$ ion and the principal residues of the IvG stabilizing the ion in the binding pocket. For E- and Z-PSL, conformers. The value for the crystal structure comprising an isolated PDS unit interacting with IgV are also given as a reference.

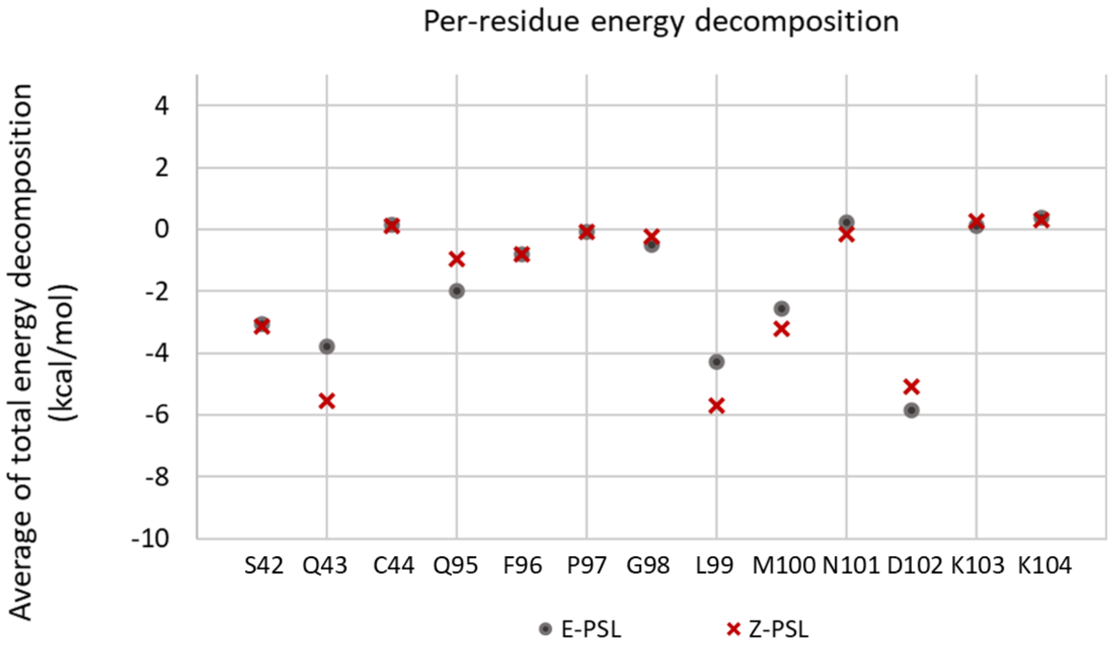


Figure S5. Per residue MMPBSA contributions.

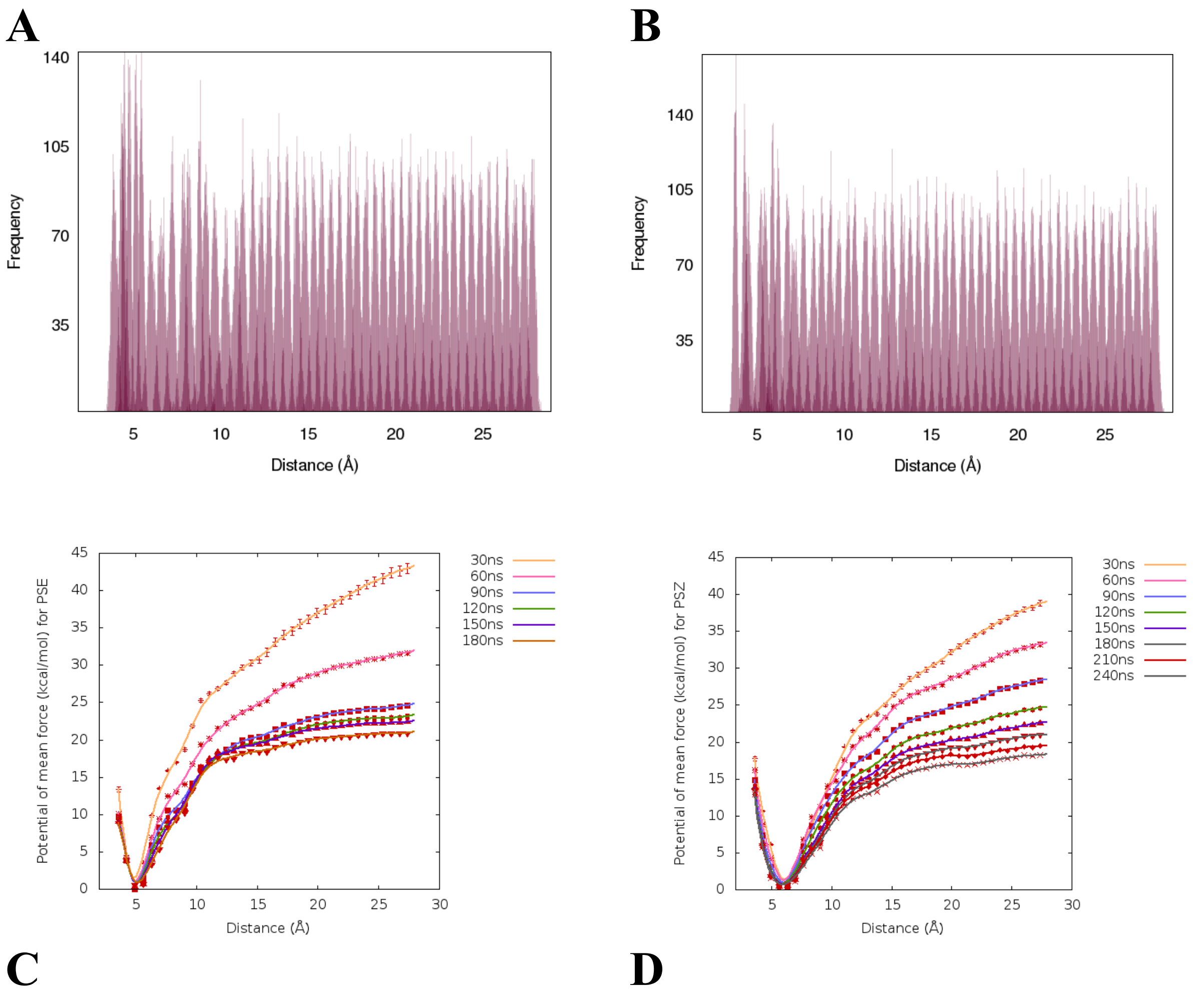


Figure S6. Distribution of the evolution of the collective variable around the different umbrella sampling for the E-PSL (A) and Z-PSL (B) conformer showing the overlap between the windows. Evaluation of the PMF convergence for E-PSL (C) and Z-PSL (D).

| Entity: | | Force constant | E | Z |
|---|---|---|---|---|
| Bonds (nm): | NN | $1.40 \times 10^7$ | 0.1270 | 0.1255 |
| | CN | $0.72 \times 10^7$ | 0.1425 | 0.1440 |
| Angles (deg): | CNN | 650.0 | 114.0 | 119.0 |
| | CCN | 560.0 | 120.0 | 120.0 |
| Dihedrals (deg): | CNNC | 70.0 | 180 | 180 |
| | CCNN | 6.0 | 180 | 180 |

Table S2. Force field parameters used to enforce the correct geometry around the isomerizable CNNC bond as taken by Osella et al. (J. Phys. Chem. C 2020, 124, 8310–8322).
